# *DESpace2*: detection of differential spatial patterns in spatial omics data

**DOI:** 10.1101/2025.06.30.662268

**Authors:** Peiying Cai, Mark D. Robinson, Simone Tiberi

## Abstract

Spatially resolved transcriptomics (SRT) enables the investigation of mRNA expression in a spatial context. While several SRT analysis frameworks have been developed, the vast majority of them focus on analyzing individual samples, and do not allow for comparisons of spatial gene expression patterns across experimental conditions, such as healthy *vs*. diseased states. Here, we present an approach to identify so-called differential spatial patterns (DSP), i.e., genes that exhibit changes in spatial expression between groups of samples across conditions. Our framework processes diverse SRT data types and detects DSP by performing differential gene expression testing across conditions. Notably, this comparison is not currently available in any other spatial omics method. In addition to detecting DSP across conditions, our framework includes two key features. First, it can identify tissue regions where expression changes across conditions. Second, it can detect spatial gene expression pattern changes across more than two experimental conditions, with flexible models. With *ad hoc* simulations, we demonstrate that our approach has good true positive rates and well-calibrated false discovery rates. Applied to experimental data, our method identifies biologically relevant DSP genes while maintaining computational efficiency. Our framework has been implemented within the *DESpace* Bioconductor R package (from version 2.0.0).

## 1 Introduction

Spatially resolved transcriptomics (SRT) technologies measure gene expression levels while preserving the tissue spatial context, within the local aggregate of multiple cells (termed spots), at the single-cell level, or even at subcellular level. This enables the investigation of spatial gene expression patterns [34,39,41], uncovers cellular heterogeneity [14], facilitates neighborhood analysis [11], enhances our understanding of disease mechanisms [6, 12, 48], and even allows for RNA expression analysis at the subcellular level [4].

To fully leverage the potential of SRT data, several analytical methods have been developed. Among these, two standard steps are the identification of spatial domains (also known as spatial clustering), and the detection of spatially variable genes (SVGs). While single-cell RNA sequencing data allows for cell grouping based on transcriptional profiles, SRT data clustering integrates both mRNA information and spatial context to identify spatial regions of coherent gene expression within a tissue. To date, a wide range of spatially-aware clustering methods have been developed for this purpose, including *BayesSpace* [47], *stLearn* [26], *Banksy* [35], *BASS* [15], *STAGATE* [10], and *GraphST* [18]. These methods utilize diverse modeling strategies, such as graph-based deep learning [10, 18] or Bayesian frameworks [15, 47]. While most are designed for single-sample analysis, some can also perform joint clustering across multiple samples [10, 35].

Methods for identifying highly variable genes (HVGs) and differentially expressed genes (DEGs), originally developed for bulk and single-cell transcriptomics, are commonly used for feature selection and cluster characterization [2, 19, 31]. However, these methods cannot explicitly incorporate spatial information. In contrast, SVGs methods [3, 40, 44] leverage spatial coordinates to identify genes with non-uniform expression patterns across the tissue. These spatially-aware approaches provide an alternative to traditional feature selection tools, and can help identify potentially biologically-informative genes for further experimental validation.

Most tools for SRT data can only work with individual samples; among the few exceptions, *DESpace* [3] supports biological replicates, while *SPADE* [29] allows comparisons between experimental conditions. Nonetheless, the original version of *DESpace* [3] does not compare groups of samples, while *SPADE* [29] is restricted to comparing two individual samples only, and has high computational demands, which makes it impractical for large-scale, multi-sample, multi-condition datasets. As a result, no method can currently compare, from SRT data, spatial patterns across groups of samples.

Here, we introduce a framework to compare spatial patterns from multi-sample, multi-condition SRT data, and identify differential spatial pattern (DSP) genes, i.e., genes whose spatial expression profiles vary between two or more experimental conditions (Figure 1). A DSP gene may, for example, be spatially variable (SV) in one condition but not in another one, or, alternatively, it may be SV in all conditions but with different spatial patterns. In contrast, genes with uniform abundance across space in all conditions, and SVGs with unchanged spatial structure between conditions, are not considered DSPs. Identifying DSP genes can provide valuable insight into how gene expression pattern vary across experimental conditions, facilitating the understanding of the underlying biological processes.

**Figure 1.**
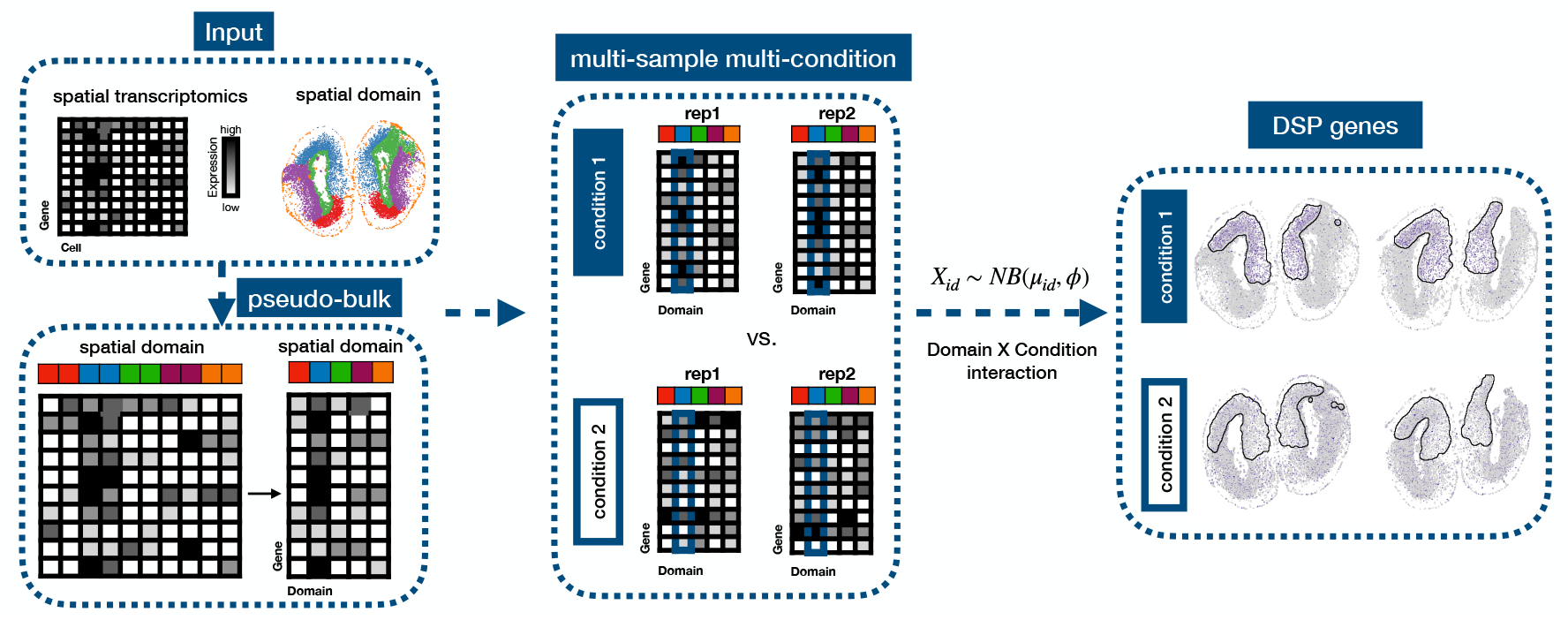
Overview of the *DESpace2* framework for DSP analysis. Given spatial transcriptomics data and spatial domain annotations, the workflow first performs pseudo-bulking and then conducts differential expression analyses across multiple samples and experimental conditions to identify DSP genes; in the example shown (right panel), a gene has a differential level of expression in one condition versus the other within the same domain.

## 2 Materials and methods

### 2.1 Overview

To detect DSP genes, we expanded our previous framework, *DESpace* [3], which identifies SVGs using pre-annotated spatial clusters, and fits *edgeR* [31] negative binomial model to the mRNA abundance of each spot, either a single cell or a multi-cellular bin, with spatial clusters as covariates. In this work, we still employ the *edgeR* negative binomial model, but use a more complex design, that includes (at least) three covariate sets: spatial domains, conditions, and their interactions. These coefficients capture the average spatial structure of gene expression, condition-specific changes in abundance, and how spatial patterns change between conditions, respectively. To make comparisons between the spatial (gene expression) structures of multiple conditions (i.e., DSP), we test whether the interaction terms between conditions and spatial domains differ from zero. Two hypothesis testing frameworks are available: the quasi-likelihood F-test (QLF) and the likelihood ratio test (LRT). By default, our package uses the QLF framework due to its improved error control, because it captures the dependency of the observations coming from the different clusters associated to the same individual.

Our two-step framework shares similarities with DEG and marker gene methods, that, although not originally designed for detecting DSP, can be adapted to perform that task. In particular, the *spatialLIBD* R package [25] computes pseudo-bulk counts in each spatial cluster (i.e., adding gene counts from all spots/cells in a cluster), and performs differential testing between clusters using *limma* [30], based on pairwise t-tests or ANOVA F-statistics. To detect DSP, the same strategy can be adapted by testing, in each individual cluster, for differences between experimental conditions, and aggregating results across all clusters (see Supplementary Details). Similarly, marker gene methods, such as *scran*’s `findMarkers` [20] and *Seurat* ‘s `FindMarkers` [33], designed to compare clusters in individual samples, can also be used to detect DSP by comparing groups of samples in individual clusters, and again aggregating results across clusters (see Supplementary Details). However, marker methods treat each spot as an independent replicate, neglecting the dependency between spots from the same sample, which makes them prone to inflated false positive rates [13, 33, 36, 49].

Although our framework is conceptually related to marker gene methods and *spatialLIBD*, it has a key difference: rather than testing for changes within individual clusters individually (i.e., one test per cluster), we evaluate overall changes across all spatial clusters, capturing global spatial effects in a single test. In addition, our framework can also reveal the specific areas of the tissue affected by DSP, identifying the spatial pattern that displays the biggest difference between conditions. Furthermore, our tool can also compare more than two conditions, and the evolution of spatial patterns across time. Finally, because of its pseudo-bulk structure, our approach is computationally efficient and is thus compatible with any type of SRT data. As our current approach has been implemented within the pre-existing *DESpace* Bioconductor/R package (from version 2.0.0); below, we will refer to our DSP framework simply as *DESpace2*.

### 2.2 Inference

Consider a SRT dataset with *N* total samples, collected under *N*_*c*_ experimental conditions, where each tissue section has been partitioned into *D* spatial domains, and where the *d*-th domain is associated to *N*_*d*_ spots. Within each sample, we sum the abundance of all spots belonging to a given domain to compute pseudo-bulk abundances, i.e., total abundance of a sample within a domain. For a given gene (index omitted for clarity), let *X*_*id*_ denote the pseudo-bulk abundance in sample *i* and domain *d*; we assume that *X*_*id*_ is a negative binomial random variable:

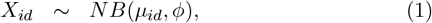

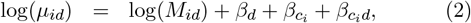

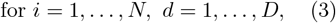

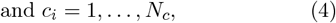

where *NB*(*µ, ϕ*) indicates a negative binomial distribution with mean *µ* and variance *µ*(1 + *µϕ*); *ϕ* is the gene-specific dispersion; *M*_*id*_ denotes the effective library size for sample *i* in domain *d*, calculated as the domain-specific total count multiplied by the trimmed mean of M values (TMM) normalization factor [32]; *β*_*d*_ represents the coefficient for spatial domain 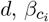 is the coefficient for condition *c*_*i*_ to which sample *i* belongs, and 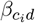 captures the interaction between domain *d* and condition *c*_*i*_.

We use a global test, to determine whether the spatial structure of gene expression varies between conditions; in particular, we test the null hypothesis that all interaction coefficients are zero:

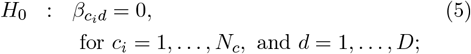

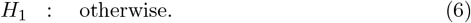

This system of hypotheses can be tested using either a likelihood ratio test (LRT) [46] or a quasi-likelihood F-test (QLF) [7, 21]. Both tests use the same negative binomial generalized linear model defined in (1), but differ in how coefficient uncertainty is quantified. The LRT is based on the negative binomial log-likelihood with fixed dispersion *ϕ*, which often results in a more sensitive but potentially liberal test. In contrast, the QLF framework uses gene-specific quasi-likelihood dispersions together with empirical Bayes shrinkage, modeling the association among the multiple observations (one per cluster) from the same individual. As a result, the QLF test provides more robust error control and better calibrated false discovery rates.

We also employ an individual-domain test to pinpoint the main DSP region, i.e., the domain where spatial patterns vary the most across conditions. To this aim, we test the interaction terms of each spatial domain; e.g., for domain *d*, we test the following system of hypotheses:

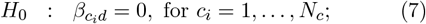

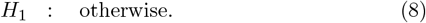

This test can be performed via a QLF or LRT approach. In all analyses, model fitting and differential testing use *edgeR* [8, 23, 31], while Benjamini-Hochberg p-value correction [1] is applied to account for multiple testing, and control for the false discovery rate (FDR).

### 2.3 Simulation studies - anchor data

To assess the performance of our approach, we designed benchmarks utilizing real and semi-simulated data, with all analyses based on one of two SRT datasets, referred to as *LIBD* [22], and *ARTISTA* [45].

The *LIBD* [22] dataset, obtained from the Visium platform [37], consists of 12 samples from three human brain donors. Pathologists manually annotated all samples, separating the tissue into white matter and up to six cortical layers (Supplementary Figure 1). Each sample of this dataset contains, on average, 33,538 genes and 3,973 spots; after filtering (see Supplementary Details), we analyzed 14,628 genes across 3,936 spots (Supplementary Table 1). The number of genes shared across samples was 33,538 before filtering and 13,743 after filtering. The *ARTISTA* [45] dataset, obtained from Stereo-seq [5] technology, captures axolotl telencephalon regeneration at single-cell resolution. The study aims to study brain healing after injury, and includes tissue samples from five distinct regeneration stages (days post-injury: DPI), with three to four sections collected from each stage, resulting in 16 samples in total. In the absence of pre-annotated spatial patterns, we applied *Banksy* multi-sample clustering to group cells into five distinct spatial clusters, which are consistent across samples (Supplementary Figure 2). On average, each sample includes measurements for 27,077 genes across 9,389 cells (Supplementary Table 1). After filtering (see Supplementary Details), we retained 13,890 genes common to all samples and an average of 9,215 cells per sample.

### 2.4 Simulation studies - SV profiles

In order to generate realistic data, we used the *LIBD* and *ARTISTA* datasets as anchors to generate several *semi-simulated* datasets. In each simulated dataset, we allocated replicate samples to two artificial conditions. Specifically, for the *LIBD* data, one group includes a slice from three samples, while the other condition includes a different slice from the same three donors. For the *ARTISTA* data, we used eight tissue sections from four regeneration stages (2, 5, 10, and 20 DPI), and randomly assigned one section of each stage to one condition, and used the remaining section for the other condition.

Using pre-annotated spatial domains (i.e., pathologist annotations for the *LIBD* data, and *Bansky* clusters for the *ARTISTA* data; see Supplementary Figures 1-2), we generated a variety of spatial patterns by re-allocating real measurements across the tissue in different ways (see Supplementary Details). In particular, we obtained uniform as well as SV patterns that correspond to real structures, e.g., the brain cortex layers in the *LIBD* dataset. Uniform structures were generated by randomly permuting observation across the entire tissue, while spatial structures were created by randomly selecting a spatial domain to have higher (main text results) or lower (shown in Supplementary Figures) abundance compared to the rest of the tissue (Supplementary Figures 3-4).

In this way, we generated four distinct scenarios - *DSP1, DSP2, NULL1*, and *NULL2* (Figure 2) - and randomly assigned genes to each in equal proportions (25% per scenario). In the DSP scenarios, gene expression patterns differ between conditions: *DSP1* genes are highly (or lowly) abundant in a randomly selected spatial domain in one condition (selected at random), and have uniform expression in the other condition; *DSP2* genes are SVGs in both conditions but with different spatial expression structures in the two conditions (two distinct randomly selected spatial domains). In contrast, the NULL scenarios have identical spatial structures in both conditions: *NULL1* genes are spatially uniformly expressed in both conditions, while *NULL2* genes are SVGs sharing the same spatial pattern in both conditions (again, with a randomly selected spatial domain). The consistent spatial structure of *NULL2* genes in the same clusters across samples facilitates accurate detection of spatial domains when combining the four distinct scenarios.

**Figure 2.**
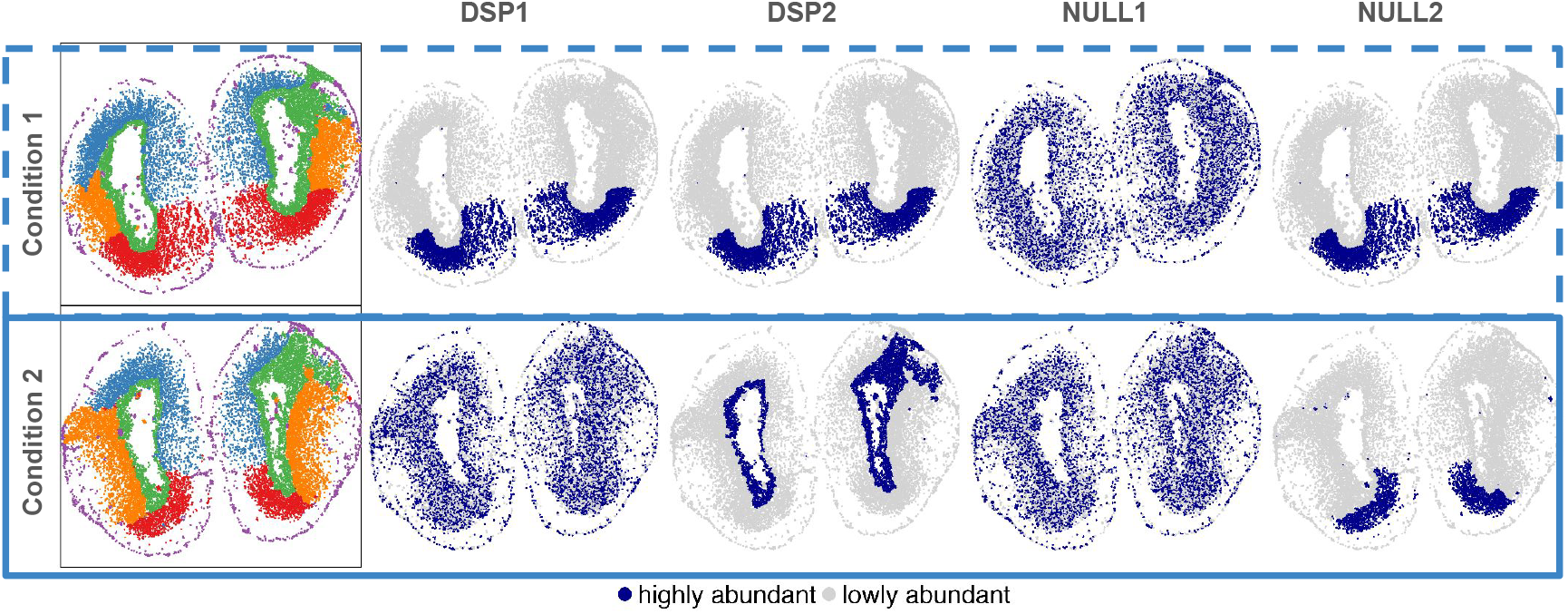
Simulation design of the four simulated categories: *DSP1, DSP2, NULL1*, and *NULL2* using the *ARTISTA* dataset. The first column shows *Banksy* spatial domains, and the next four columns present example of simulations for each category. Two rows correspond to different conditions.

## 3 Results

### 3.1 Simulation studies - global test

In the *LIBD* and *ARTISTA* simulated dataset, we re-estimated the spatial patterns via *BayesSpace* and *Bansky*, respectively, leading to five consistent spatial domains across samples in both datasets (see Supplementary Details). In our benchmarks, all methods used these (noisy) estimated clusters, and not the original domains used to generate the simulated data. The simulated data was analyzed via *DESpace2* (DSP functionality with QLF and LRT testing), *scran*’s `findMarkers` [20], *Seurat* ‘s `FindMarkers` [33] (both in the single-cell and pseudo-bulk version), and *spatialLIBD’s wrapper* [25]. Since no existing approach is specifically designed to detect changes in the spatial structure of gene expression, between groups of samples, we adapted these methods to identify DSP genes. We excluded *SPADE* [29] from our analyses because it is designed to only compare individual samples, and has an extreme computational cost: in our tests, it required approximately two hours to fit a single gene with approximately 4,000 spots.

The left panels of Figure 3 display the true positive rate (TPR) *vs*. false discovery rate (FDR) for all methods in our simulations, which include all four scenarios (i.e., *DSP1, DSP2, NULL1* and *NULL2*) combined. Across our semi-simulated benchmarks, *DESpace2* controls the false discovery rate and offers competitive statistical power. `findMarkers` and `FindMarkers` also achieve a good TPR, but exhibit an inflated FDR, leading to a significant number of false discoveries. The inflation of false positives has been previously reported in the literature [13, 33, 36, 49], and is possibly due to the fact that marker methods treat each cell/spot as an independent replicate, and neglect the dependency among cells/spots from the same sample. Conversely, pseudobulk-based `FindMarkers` and *spatialLIBD*, control the FDR, but have lower statistical power, possibly due to the fact that they compare pairs of clusters rather than jointly capturing global changes across all clusters, as *DESpace2* does.

**Figure 3.**
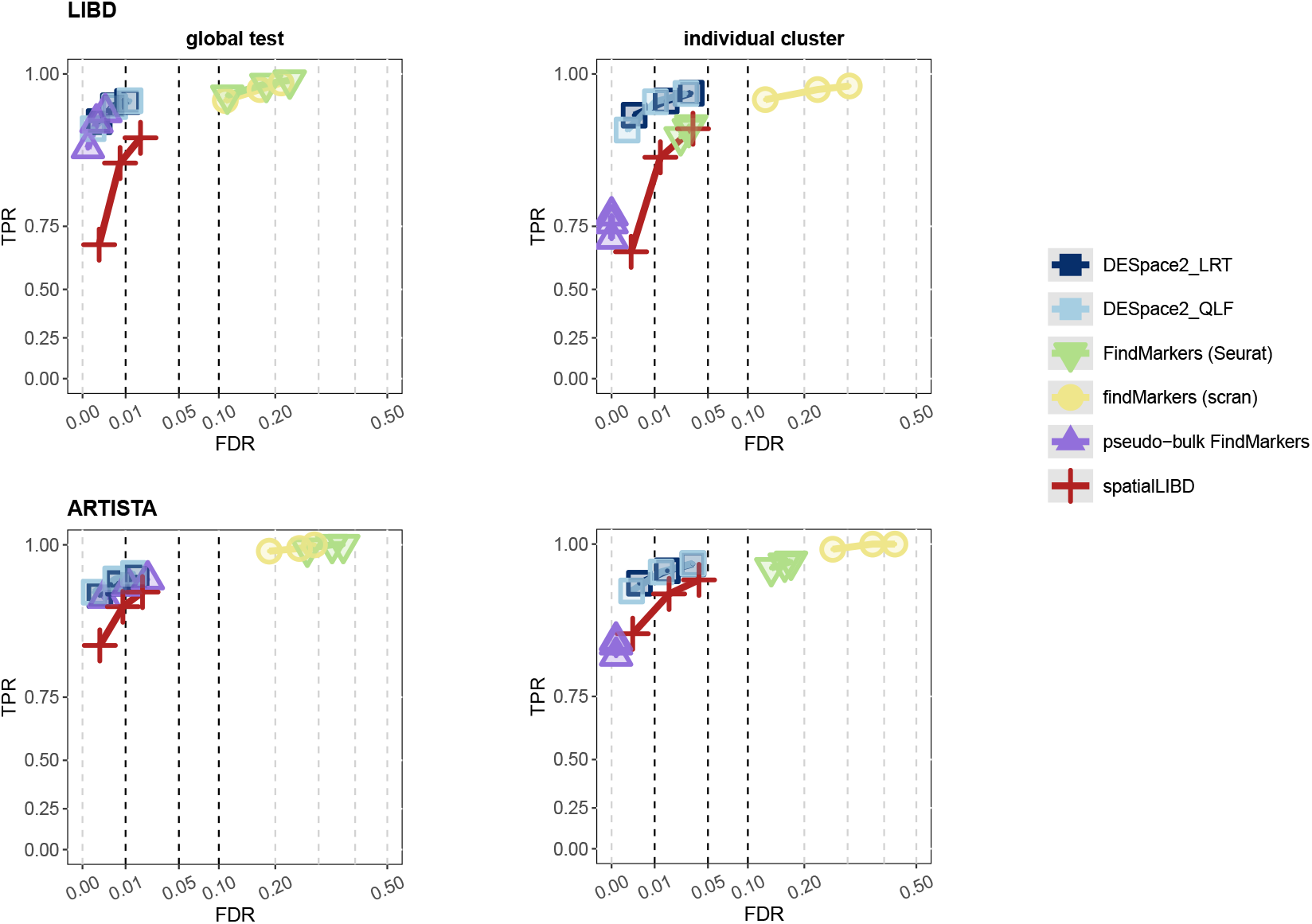
TPR *vs*. FDR for DSP gene detections in our simulations. Top and bottom rows refer to the *LIBD* and *ARTISTA* data, respectively, while left and right columns indicate the global and individual-domain tests, respectively.

We also studied the impact of clustering by comparing *DESpace2* results obtained on *BayesSpace*/*Banksy* clusters with those obtained using the original patterns the data was simulated from (i.e., the ground truth): we found that the noise arising from the estimated patterns have a small impact on results (Supplementary Figure 5), likely because the recomputed clusters were very similar to the original annotation used in simulation.

Additionally, we investigated false positive rates; to this aim, we analyzed the distribution of raw p-values for NULL1 and NULL2 genes (i.e., genes with uniform expression or identical spatial patterns across conditions). In all scenarios, *DESpace2* shows no inflation of false positives, while all our competitors display inflated false positive rates (Supplementary Figure 7); this is likely because our competitors perform a test for each spatial pattern, and the global score is obtained as the minimum of the individual cluster p-values. Indeed, as expected, the individual-domain p-values are more conservative (see next Section).

Supplementary Figure 6 presents results from the simulation where the selected spatial domain has lower abundance compared to the rest of the tissue. Here, the spatial signal is diluted because the high abundance region (represented by *D* − 1 domains) is wide, and this makes DSP detection more complex. Results are similar to those shown above (with lower TPRs for all methods), and the ranking of methods is stable.

### 3.2 Simulation studies - individual-domain

We also evaluated *DESpace2*’s ability to identify the domains where gene abundance varies across conditions. The individual-domain test requires at least three clusters (with two it is equivalent to the global test), and may be particularly useful when analyzing datasets with many clusters, as it may help biologists to focus on specific domains of interest.

To validate this framework, we considered the same simulation used in the global test above, and aimed to identify the main pattern that varies between conditions. *DSP1* genes are SV in one condition only: the key DSP cluster is the one selected to have higher or lower abundance in the SV condition. *DSP2* genes, genes are SVGs under both conditions, but with different spatial structures; in this case, we have two main DSP clusters are the two clusters with higher or lower abundance in each condition (Figure 2).

For each method, we tested all domains, and counted how often the most significant domain(s) corresponds to the main DSP cluster (for *DSP1* genes), or the main two DSP clusters (for *DSP2* genes). For *DSP1* genes, *DESpace2* identified the main DSP cluster in 96-100% of genes; `FindMarkers` and `findMarkers` lead to similar percentages, while *pseudo-bulk* `FindMarkers` and *spatial-LIBD* displayed lower accuracy. For *DSP2* genes, which generally yields higher scores than *DSP1, DESpace2* using the QLF identified the two main DSP clusters in 99.9 (96.0) and 100 (98.4)% of genes in our semi-simulated *LIBD* and *ARTISTA* dataset, respectively, when the main clusters had higher or lower abundance, with competitors showing comparable performance (Table 1 and Supplementary Table 2).

We also generated TPR *vs*. FDR plots (right panel of Figure 3). In this case, in DSP genes all domains are DSP, even if most with minor changes between conditions. Therefore, in DSP genes, we only considered the main spatial clusters involved (i.e., the main cluster for *DSP1*, and main two for *DSP2*); instead, as controls we included all clusters from NULL patterns. Similarly to the global test, *DESpace2* has good FDR control, and higher TPR than competitors. Notably, the *DESpace2* LRT test achieves higher sensitivity than its QLF counterpart under this setting.

We also plotted the densities of NULL1 and NULL2 p-values (Supplementary Figures 8-9): all methods provide approximately uniform (or slightly conservative) null p-values, except for `FindMarkers` and `findMarkers` where a general inflation towards 0 is observed, indicating increased false positive rates.

In the simulation where the main spatial domain is characterized by lower abundance (Supplementary Figure 4), results are similar, with lower statistical power for all methods given by the more subtle DSP structure (Supplementary Figure 6).

### 3.3 Simulation studies - sensitivity analyses

Given that our framework relies on predefined spatial domains, we further assessed each method’s robustness under two conditions: mis-clustering and clustering resolution. For mis-clustering, two scenarios were examined. First, a random subset of spots (1%, 5%, 10%, and 20%) was randomly reassigned to incorrect clusters. Second, we simulated datasets using five spatial domains, and forced *Bansky* to recover an incorrect number of clusters (varying from three to ten).

As the proportion of mis-assigned cells increases, all methods show reduced TPR (Figure 4 and Supplementary Figure 10); nonetheless, *DESpace2* consistently achieves higher TPRs than competing approaches, while controlling the FDR in both global and individual-domain tests. Similarly, when the number of clusters was mis-specified, all methods exhibit reduced performance, particularly when the number of clusters was underestimated (Supplementary Figure 11). Instead, when the number of sub-clusters is overestimated, all methods show smaller declines in TPR; importantly, across the evaluated scenarios, *DESpace2* maintains a well calibrated FDR. In this setting, the QLF version of *DESpace2* is more stable than the LRT version; this is consistent with QLF’s use of quasi-likelihood dispersion estimation, which captures intra-sample variability more effectively, and therefore tends to be more robust when the model is misspecified.

**Figure 4.**
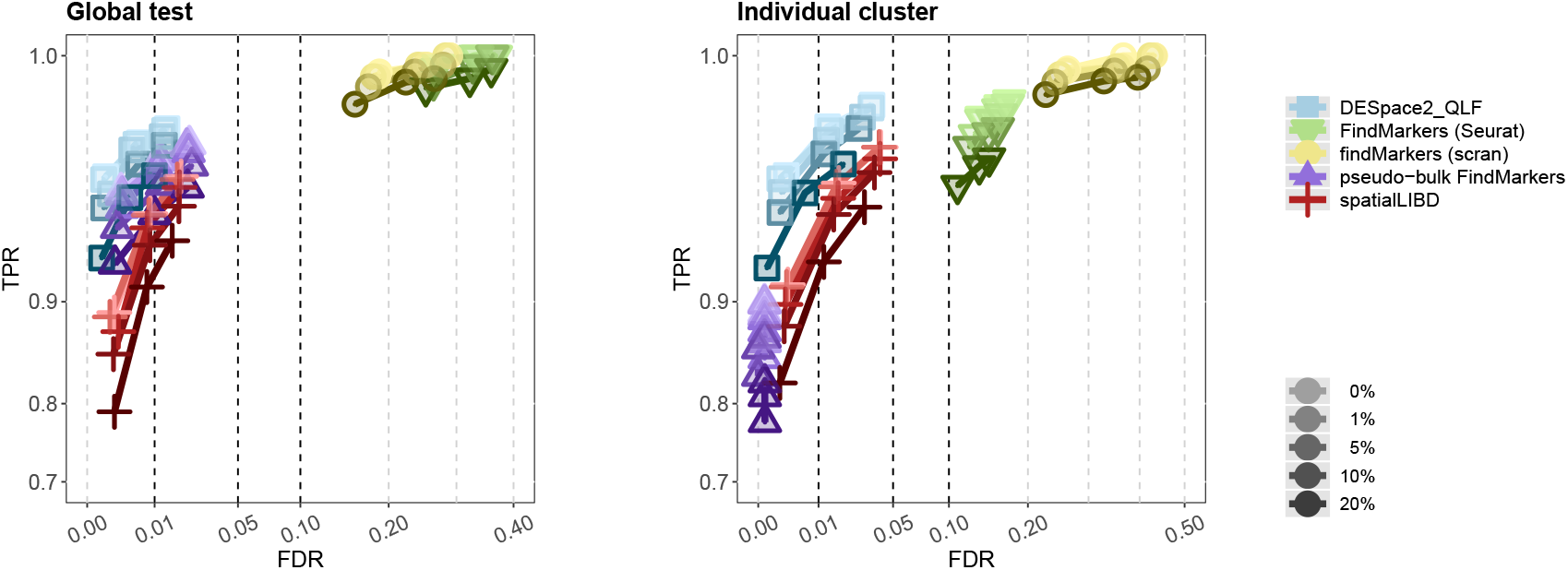
TPR *vs*. FDR for DSP gene detection in sensitivity analyses using the *ARTISTA* dataset. A subset of cells (0%, 1%, 5%, 10%, and 20%) was randomly reassigned to incorrect clusters to simulate varying levels of mis-clustering. Right and left panels refer to the global and individual-domain tests, respectively.

Next, we evaluated robustness to clustering resolution by simulating datasets with varying numbers of clusters (Supplementary Figures 12-13). In the individual test, *DESpace2* consistently achieves higher TPRs than competitors, particularly with fewer clusters (Supplementary Figure 13). Clustering resolution has a marginal impact on the FDR for both the QLF and LRT implementations. In the global test, *DESpace2_QLF* has the highest TPR with two, four, and six clusters, and remains comparable to the pseudobulk-based `FindMarkers` at higher resolutions (Supplementary Figure 12). The LRT version follows a similar trend at low resolutions but shows reduced detection power and a marginal increase in FDR as the number of clusters grows. This trend likely reflects the increasing number of coefficients that the LRT needs to estimate in the global test as the clustering resolution increases, making it sensitive to model complexity. Overall, these results highlight the robustness of *DESpace2*, particularly under the QLF-based test, which, in our benchmarks, shows well-calibrated error rates while maintaining good power.

### 3.4 Applications to real data

We applied all methods to the original *ARTISTA* dataset, comparing two conditions (2 and 20 DPI), which are the earliest and latest time points to have replicates, and may possibly represent the most distinct two groups to analyze. Additionally, for *DESpace2* and *SpatialLIBD*, which both support comparisons across more than two conditions, we also included all five time points (2, 5, 10, 15, and 20 DPI), each with three or four samples (Supplementary Table 1). In absence of manually annotated clusters, spatial clusters were computed using *Banksy*, which was jointly applied to all 16 samples across five time points. For the two-group comparison, the data were then subset to 2 and 20 DPI, ensuring that both two- and five-group comparisons used the same spatial domain annotations and number of genes. For the two-condition analysis, *DESpace2* and *spatialLIBD* identified a similar number of significant genes (around 250), while `FindMarkers` and `findMarkers` reported over 7,800 significant genes, out of 13,890, which may reflect the inflation of false positives observed in simulation studies (Table 2).

**Table 1.** Individual cluster results. Percentage of times the method identified the main SV cluster in *DSP1* and when the top two detected domains both corresponded to the main SV clusters in *DSP2*, using *BayesSpace* for *LIBD* and *Banksy* for *ARTISTA*.

|  | Method | *LIBD* | *ARTISTA* |
| --- | --- | --- | --- |
| *DSP1* | *DESpace2_LRT* | 98.1 | 99.3 |
|  | *DESpace2_QLF* | 96.6 | 99.5 |
|  | *Seurat's FindMarkers* | 94.4 | 98.8 |
|  | *scran's findMarkers* | 94.1 | 98.8 |
|  | pseudobulk-based **Findmarkers** | 85.3 | 88.9 |
|  | *spatialLIBD* | 71.5 | 80.4 |
| *DSP2* | *DESpace2_LRT* | 100 | 100 |
|  | *DESpace2_QLF* | 99.9 | 100 |
|  | *Seurat's FindMarkers* | 99.7 | 100 |
|  | *scran's findMarkers* | 99.7 | 100 |
|  | pseudobulk-based **Findmarkers** | 99.6 | 99.6 |
|  | *spatialLIBD* | 96.2 | 99.7 |

**Table 2.** Summary of genes detected by each method in the *ARTISTA* dataset. The table shows the number of genes detected, among the top 200 results returned by each method. Gene sets were retrieved from the Molecular Signatures Database (MSigDB v2024.1.Hs) for the following keywords: “Wound healing” and “Regeneration”. These sets include 434 and 203 genes, respectively, of which 127 and 257 were present in our dataset. “Overall” represents the union of both sets (603 genes total; 366 retained). The table also reports the average ranking of the genes in each list (lower values indicate higher significance), and the total number of genes called significant by each method.

| Method | Identified in top 200 |  |  | Average ranking |  | # of genes |
| --- | --- | --- | --- | --- | --- | --- |
|  | Regeneration | Wound healing | Overall | Regeneration | Wound healing |  |
| *DESpace2_QLF*<br>(5 groups) | 5 | 15 | 20 | 2629 | 2914 | 848 |
| *DESpace2_QLF* | 4 | 12 | 16 | 1458 | 1599 | 71 |
| *DESpace2_QLF*<br>(individual test) | 5 | 10 | 15 | 4386 | 4613 | 23 |
| *DESpace2_QLF*<br>(individual test; 5 groups) | 3 | 10 | 13 | 4765 | 5371 | 372 |
| *Seurat's FindMarkers* | 7 | 6 | 13 | 4054 | 5100 | 7809 |
| *spatialLIBD* | 7 | 6 | 13 | 5865 | 6675 | 289 |
| spline-based *DESpace2_QLF*<br>(5 groups) | 5 | 6 | 11 | 1402 | 1534 | 130 |
| *scran's findMarkers* | 6 | 5 | 11 | 4122 | 5159 | 7016 |
| pseudobulk-based<br>*Seurat's FindMarkers* | 6 | 5 | 11 | 5277 | 6176 | 2957 |
| *scran's findMarkers*<br>(5 groups) | 5 | 5 | 10 | 3703 | 4691 | 8732 |
| *spatialLIBD*<br>(5 groups) | 0 | 2 | 2 | 5994 | 6526 | 6544 |

Unlike simulations, real data does not have a ground truth; to overcome this, we generated two lists of potentially relevant genes by retrieving the gene sets “GOBP_WOUND_HEALING” and “GOBP_REGENERATION” from the Molecular Signatures Database (MSigDB v2024.1.Hs) [16, 17, 38]. Because axolotl-specific functional curation is limited, these human-curated MSigDB sets served as proxies for assessing biological relevance. We then calculated the top 200 (i.e., most significant) genes from each method, and counted how many of those appear in the two lists. Additionally, for each method, we computed the average ranking of the genes in the wound healing and regeneration lists, where a ranking of 1 indicates the most significant gene returned by a method. The top 200 genes from *DESpace2*, in both its 2-condition and 5-condition comparisons, contains more genes from the wound healing and regeneration lists than competitors; furthermore, those genes also have a smaller average ranking (i.e., more significant) (Table 2; LRT-based results shown in Supplementary Table 3). We also applied *DESpace2* with a spline model to account for the effect of time (i.e., spline-based *DESpace2*), which, despite detecting fewer interesting genes among the top 200, achieves a better (lower) average ranking.

Furthermore, we retrieved genes related to healing and regeneration using data from The Human Protein Atlas website [27]. The performance of methods is consistent, with *DESpace2* detecting either more or the same number of healing- and regeneration-related genes as competing methods, while ranking higher (i.e., with a lower average ranking) (Supplementary Table 4).

Finally, we investigated the computational efficiency of each method by measuring their runtime on the *ARTISTA* dataset (Figure 5 and Supplementary Figure 14). As expected, *spatialLIBD, DESpace2*, and pseudobulk-based `FindMarkers`, allow of which use pseudobulk counts, have a much lower runtime compared to *scran*’s `findMarkers` and spot-level *Seurat*’s `FindMarkers`, which use single spot/cell data. Notably, *Seurat* ‘s `FindMarkers` is substantially slower than *scran*’s `findMarkers`, as previosusly reported [28]. Importantly, for *DESpace2*, we report the costs for performing global test only (i.e., *DESpace2_QLF_global*; 14 seconds), and for running individual-pattern tests (i.e., *DESpace2_QLF_individual*; 36 seconds); the runtimes for *DESpace2_LRT* are nearly identical (14 and 38 seconds for the global and individual tests, respectively). Note that all methods in this analysis rely on precomputed spatial clusters. As such, we do not report for the computational cost of generating these domain annotations, and only consider the runtime of each method. The runtime of *Bansky* spatial clustering was 97 minutes for the 5-group comparison (16 samples).

**Figure 5.**
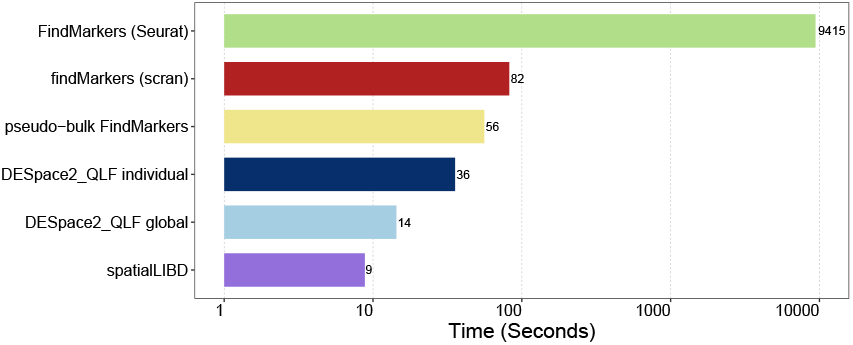
Runtime (in seconds) for each method on the *ARTISTA* dataset using a 2-condition comparison (2 and 20 days post-injury) across six samples.

## 4 Discussion

We presented a novel framework, based on spatial domains and DEG testing, to identify DSP genes, i.e., genes where structure of spatial expression varies between groups of samples, which represents a type of analysis that is absent from the current spatial omics toolbox. We benchmarked our approach against diverse alternatives (e.g., adapting marker and DE methods) in *ad hoc* semi-simulated and real data analyses, where our framework demonstrates good statistical power (i.e., TPR), controls for the FDR, and is computationally efficient. Our method can detect differences across the entire tissue (i.e., global test), or test each specific cluster (i.e., individual-pattern test), identifying the main region that varies across conditions. Our tool inputs flexible user-customizable design matrices; this allows users, for instance, to compare more than two conditions, and to model time courses (e.g., via splines). Users can choose between the QLF- and LRT-based testing options, depending on their needs: QLF generally provides more stable error control and is used as the package default, whereas LRT can offer increased sensitivity when the spatial signal is diluted. Our DSP framework is freely distributed within the *DESpace* Bioconductor R package, which simplifies its usage and integration with other pipelines, and is accompanied by an example usage DSP vignette and *ad hoc* plotting functions. Furthermore, our approach is flexible and can be applied to any spatial transcriptomics technology.

We also acknowledge some limitations of our work. Our approach relies on predefined spatial clusters, which is generally manageable in practice: spatial clustering is a standard step in spatial transcriptomics analysis, with over 50 spatial clustering methods developed since 2020 [24]. Importantly, because of this dependence, our method is more sensitive to detecting DSP genes whose expression aligns with the spatial patterns provided, having lower statistical power to detect genes that do not follow these structures (found to be more rare in previous research [3, 44]). Additionally, domains that exclusively appear in one condition cannot be analyzed; nonetheless, these domains indicate a condition-specific region (i.e., present in some conditions and absent in others), rather than a DSP region. This is conceptually similar to what happens with differential state analyses [9, 42, 43], which target differences in abundance only in cell clusters that appear in all conditions, and ignore differences in cell cluster proportions.

In some cases, DSP could represent shifts in composition of the cell types within spatial clusters. Such changes may still be of direct interest, or could be adjusted for as covariates.

Overall, we believe that our DSP framework represents a useful instrument for investigating spatial omics data; additionally, it also represents an initial step for comparing groups of samples from SRT data, and may pave the road for other future methods in the field.

## Availability

Our DSP framework is freely implemented within *DESpace* Bioconductor R package (from version 2.0.0) at https://bioconductor.org/packages/DESpace; the DSP vignette is titled “Differential Spatial Pattern between conditions”, and also includes examples where time points are compared via smooth splines. The scripts and data used for the analyses presented in this study are available on GitHub (https://github.com/peicai/DESpace2_manuscript) and Zenodo (DOI: 10.5281/zenodo.17750644), respectively. The *LIBD* [22] dataset was obtained from the *spatialLIBD* R Bioconductor package, and the original *ARTISTA* [45] dataset is available at https://db.cngb.org/stomics/artista/download/. We created an ExperimentHub R Bioconductor package, called *muSpaData*, to store the processed version of the *ARTISTA* data, including the spatial clusters we computed via *Bansky*; the package is available at https://bioconductor.org/packages/muSpaData.

## Acknowledgements

We thank Robinson lab members for helpful discussions and in particular David Wissel for discussions about smooth spline basis parameterizations.

## Author contributions

P.C. implemented the approach, conducted all analyses, and drafted the manuscript, contributing to the study design. M.D.R. assisted with study design and manuscript revision. S.T. conceived the method, led the study design and contributed to editing. All authors approve the article.

