## Supplementary Information for "*DESpace2*: detection of differential spatial patterns in spatial omics data"

### 1 Supplementary Details

#### 1.1 Adjusted p-values

For all methods, we controlled the false discovery rate using Benjamini–Hochberg adjusted p-values [3].

For *spatialLIBD* [9], *scran*'s `findMarkers` [6], and *Seurat*'s `FindMarkers` [11], we computed per-gene q-values using the `perGeneQValue` function from *DEXSeq* [2, 5], to summarize and correct for multiple testing across clusters.

#### 1.2 Quality control and filtering

Quality control procedures were performed according to the guidelines outlined in the online book "Orchestrating Spatially-Resolved Transcriptomics Analysis with Bioconductor" [1].

To identify low-quality spots or cells, we used the *Scuttle*'s `addPerCellQC` function [8]. Spots/cells were evaluated based on total UMI counts, the number of detected genes (i.e., genes with non-zero UMI counts), the proportion of reads mapped to mitochondrial genes, and the estimated number of cells per spot. We then applied sample-specific thresholds to filter out low-quality spots/cells.

We also removed undetected or lowly expressed genes, retaining only those expressed in at least 20 non-zero spots/cells for downstream analysis.

Finally, to enable cross-sample analysis, we excluded genes that were not present in all samples.

#### 1.3 Spatial domains used in simulation

For the *LIBD* dataset, we used pathologist-provided annotations from the original data [7] to define spatial patterns for simulation. To harmonize the number of spatial domains across samples, we reassigned "Layer1" and "Layer2" to "Layer6" in samples where they were present, yielding five common domains (Layers 3–6 and WM) for cross-sample simulation. After generating the simulated data, spatial domains were re-estimated for each sample using *BayesSpace*. Given the clear structure of cortical layers and white matter, manual alignment of spatial domains across samples was straightforward.

In contrast, the *ARTISTA* dataset lacked predefined spatial annotations. To define consistent spatial domains across samples, we applied joint clustering using *Banksy*. Simulations were then performed based on these domains. Afterward, we re-estimated the spatial domains using *Banksy*, jointly across all samples, to provide input for each method.

### 1.4 Simulating SV patterns

To generate simulated data, we first partitioned each tissue sample into two regions, $R_1$ and $R_2$, containing $n_1$ and $n_2$ spots or cells, respectively. We then adopted the `runPatternSimulation` function from *Giotto*'s R package [4] to assign gene expression patterns based on these regions. This function selects the $n_1$ most highly expressed genes and assigns them to region $R_1$ with probability $\pi$. When $\pi = 1$, the $n_1$ highest-expressing spots/cells are assigned to $R_1$, creating a strong spatial pattern. In contrast, $\pi = 0.5$ results in uniform expression across both regions.

Next, we simulated a mixture of uniformly expressed and SV genes. Uniform patterns were generated by sampling $\pi$ from a Beta distribution with a mean of 0.5 and a standard deviation of 0.025. In contrast, SVGs were modeled using a Beta distribution with a mean of 0.9 and the same standard deviation. For each SVGs, we randomly selected a spatial domain to serve as $R_1$ and assigned the most abundant expression measurements to this region, using randomly sampled $\pi$. This approach ensured that the selected domain exhibited average higher expression compared to the rest of the tissue. Using the uniform and/or spatial patterns generated for each sample under each condition, we created four categories of possible scenarios - *DSP1*, *DSP2*, *NULL1*, and *NULL2* - and randomly assigned genes to each scenario in equal proportions (25% each).

We also generated a parallel simulation in which SVGs were lowly abundant in the selected domain(s), instead of highly abundant. This allowed us to evaluate method performance under both high- and low-abundance spatial expression settings.

### 1.5 Software versions

All analyses were performed in R (version 4.4.2, with Bioconductor packages from release 3.20).

### 2 Supplementary Tables

| Data set | Sample Ids | Before Filtering | | After Filtering | |
|---|---|---|---|---|---|
| | | # of Genes | # of Spots | # of Genes | # of Spots |
| LIBD | 151507 | 33538 | 4226 | 14226 | 4172 |
| | 151508 | 33538 | 4384 | 13835 | 4333 |
| | 151509 | 33538 | 4789 | 14561 | 4738 |
| | 151510 | 33538 | 4634 | 14331 | 4606 |
| | 151669 | 33538 | 3661 | 14547 | 3647 |
| | 151670 | 33538 | 3498 | 14183 | 3473 |
| | 151671 | 33538 | 4110 | 14926 | 4079 |
| | 151672 | 33538 | 4015 | 14634 | 3989 |
| | 151673 | 33538 | 3639 | 15115 | 3604 |
| | 151674 | 33538 | 3673 | 15878 | 3634 |
| | 151675 | 33538 | 3592 | 14626 | 3556 |
| | 151676 | 33538 | 3460 | 14672 | 3405 |
| | Average | 33538 | 3973 | 14628 | 3936 |
| ARTISTA | 2DPI_1 | 27324 | 7668 | 15520 | 7544 |
| | 2DPI_2 | 28365 | 8192 | 15633 | 8054 |
| | 2DPI_3 | 28371 | 8738 | 15734 | 8602 |
| | 5DPI_1 | 28741 | 8106 | 15995 | 7947 |
| | 5DPI_2 | 29442 | 8178 | 16221 | 8023 |
| | 5DPI_3 | 29341 | 8372 | 16213 | 8220 |
| | 10DPI_1 | 27600 | 9440 | 15623 | 9253 |
| | 10DPI_2 | 28602 | 9600 | 15932 | 9407 |
| | 10DPI_3 | 27398 | 10002 | 15421 | 9806 |
| | 15DPI_1 | 23331 | 8996 | 16289 | 8817 |
| | 15DPI_2 | 23885 | 9639 | 16548 | 9444 |
| | 15DPI_3 | 22297 | 9676 | 15714 | 9482 |
| | 15DPI_4 | 23271 | 10792 | 16404 | 10575 |
| | 20DPI_1 | 28205 | 10462 | 15892 | 10284 |
| | 20DPI_2 | 28987 | 11319 | 16389 | 11118 |
| | 20DPI_3 | 28069 | 11048 | 15822 | 10856 |
| | Average | 27077 | 9389 | 15959 | 9215 |

**Supplementary Table 1:** Number of genes and spots for each sample, before and after filtering.

| | Method | *LIBD* | *ARTISTA* |
|---|---|---|---|
| *DSP1* | *DESpace2_LRT* | 97.1 | 98.4 |
| | *DESpace2_QLF* | 96.9 | 98.3 |
| | *Seurat*'s `FindMarkers` | 94.5 | 97.4 |
| | *scran*'s `findMarkers` | 94.2 | 97.6 |
| | pseudobulk-based `Findmarkers` | 94.7 | 96.3 |
| | *spatialLIBD* | 92.5 | 95.5 |
| *DSP2* | *DESpace2_LRT* | 96.8 | 98.4 |
| | *DESpace2_QLF* | 96.0 | 98.4 |
| | *Seurat*'s `FindMarkers` | 94.9 | 99.0 |
| | *scran*'s `findMarkers` | 94.5 | 98.6 |
| | pseudobulk-based `Findmarkers` | 95.8 | 97.4 |
| | *spatialLIBD* | 92.9 | 97.0 |

**Supplementary Table 2:** Individual cluster results when the main spatial cluster has lower abundance than the rest of the tissue. Percentage of times the method identified the main SV cluster in *DSP1* and when the top two detected domains both corresponded to the main SV clusters in *DSP2*, using *BayesSpace* for *LIBD* and *Banksy* for *ARTISTA*.

| Method | Identified in top 200 | | | Average ranking | | # of genes |
|---|---|---|---|---|---|---|
| | Regeneration | Wound healing | Overall | Regeneration | Wound healing | |
| *DESpace2_LRT* | 5 | 12 | 17 | 1943 | 2016 | 229 |
| *DESpace2_LRT* (5 groups) | 5 | 12 | 17 | 3185 | 3452 | 1658 |
| *DESpace2_LRT* (individual test; 5 groups) | 3 | 10 | 13 | 4979 | 5318 | 1322 |
| *DESpace2_LRT* (individual test) | 1 | 10 | 11 | 3574 | 3650 | 152 |
| spline-based *DESpace2_LRT* (5 groups) | 4 | 6 | 10 | 1588 | 1714 | 261 |

**Supplementary Table 3:** Summary of genes detected by *DESpace2* under the likelihood ratio testing framework in the *ARTISTA* dataset. The table reports the number of genes detected among the top 200 results, the corresponding MSigDB (v2024.1.Hs) "Wound healing" and "Regeneration" set overlaps, and the average ranking and total number of significant genes.

| Method | Identified in top 200 | | | Average ranking | | # of genes |
|---|---|---|---|---|---|---|
| | Regeneration | Wound healing | Overall | Regeneration | Wound healing | |
| *DESpace2_LRT* | 6 | 9 | 15 | 1898 | 2120 | 229 |
| *DESpace2_QLF* | 5 | 10 | 15 | 1435 | 1703 | 71 |
| *DESpace2_QLF* (5 groups) | 6 | 9 | 15 | 2669 | 3017 | 848 |
| *spatialLIBD* | 8 | 7 | 15 | 5949 | 6862 | 289 |
| *DESpace2_QLF* (individual test) | 6 | 8 | 14 | 4334 | 4813 | 23 |
| *DESpace2_LRT* (5 groups) | 6 | 6 | 12 | 3242 | 3598 | 1658 |
| *Seurat*'s `FindMarkers` | 7 | 5 | 12 | 4246 | 5417 | 7809 |
| *DESpace2_LRT* (individual test) | 2 | 9 | 11 | 3546 | 3709 | 152 |
| *DESpace2_LRT* (individual test; 5 groups) | 3 | 8 | 11 | 4991 | 5538 | 1322 |
| *scran*'s `findMarkers` | 6 | 4 | 10 | 4258 | 5420 | 7016 |
| pseudobulk-based *Seurat*'s `FindMarkers` | 6 | 4 | 10 | 5415 | 6434 | 2957 |
| *DESpace2_QLF* (individual test; 5 groups) | 3 | 6 | 9 | 4802 | 5557 | 372 |
| spline-based *DESpace2_QLF* (5 groups) | 5 | 4 | 9 | 1435 | 1558 | 130 |
| *scran*'s `findMarkers` (5 groups) | 5 | 4 | 9 | 3800 | 4483 | 8732 |
| spline-based *DESpace2_LRT* (5 groups) | 4 | 4 | 8 | 1626 | 1742 | 261 |
| *spatialLIBD* (5 groups) | 2 | 1 | 3 | 5972 | 6630 | 6544 |

**Supplementary Table 4:** Summary of genes detected by each method in the *ARTISTA* dataset. The table shows the number of genes detected, among the top 200 results returned by each method. Gene sets were retrieved from The Human Protein Atlas website [10] using the keywords "Healing" and "Regeneration". These sets include 243 and 230 genes, respectively, of which 159 and 147 were present in our dataset. "Overall" represents the union of both sets (445 genes in total; 290 retained). The table also reports the average ranking of genes in each list (lower values indicate higher significance), as well as the total number of genes called significant by each method.

### 3 Supplementary Figures

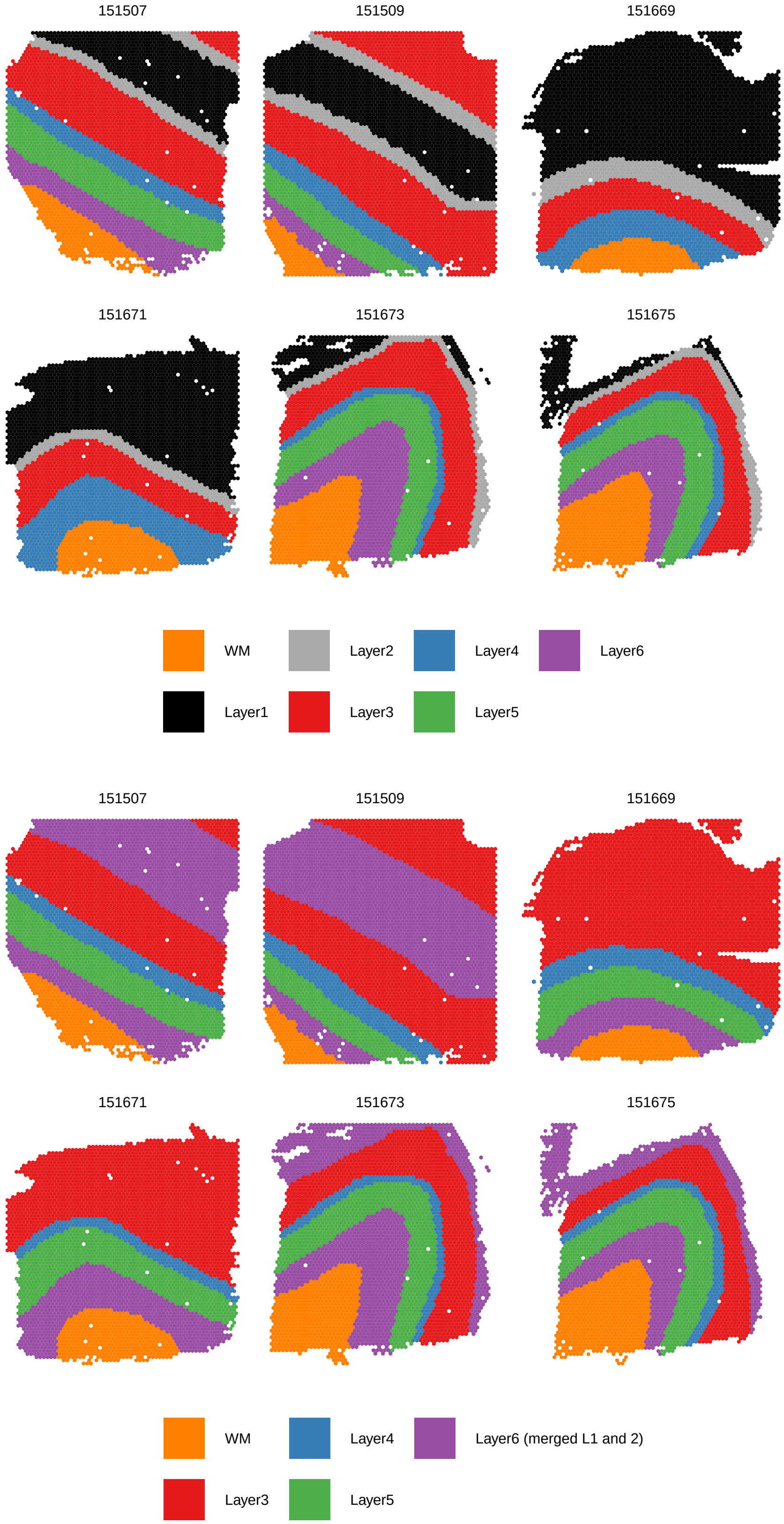


**Supplementary Figure 1:** Spatial domains for the *LIBD* dataset. Top: original pathologist annotations. Bottom: processed domains used in simulation, with Layers 1 and 2 merged into Layer 6 to harmonize domains across samples.

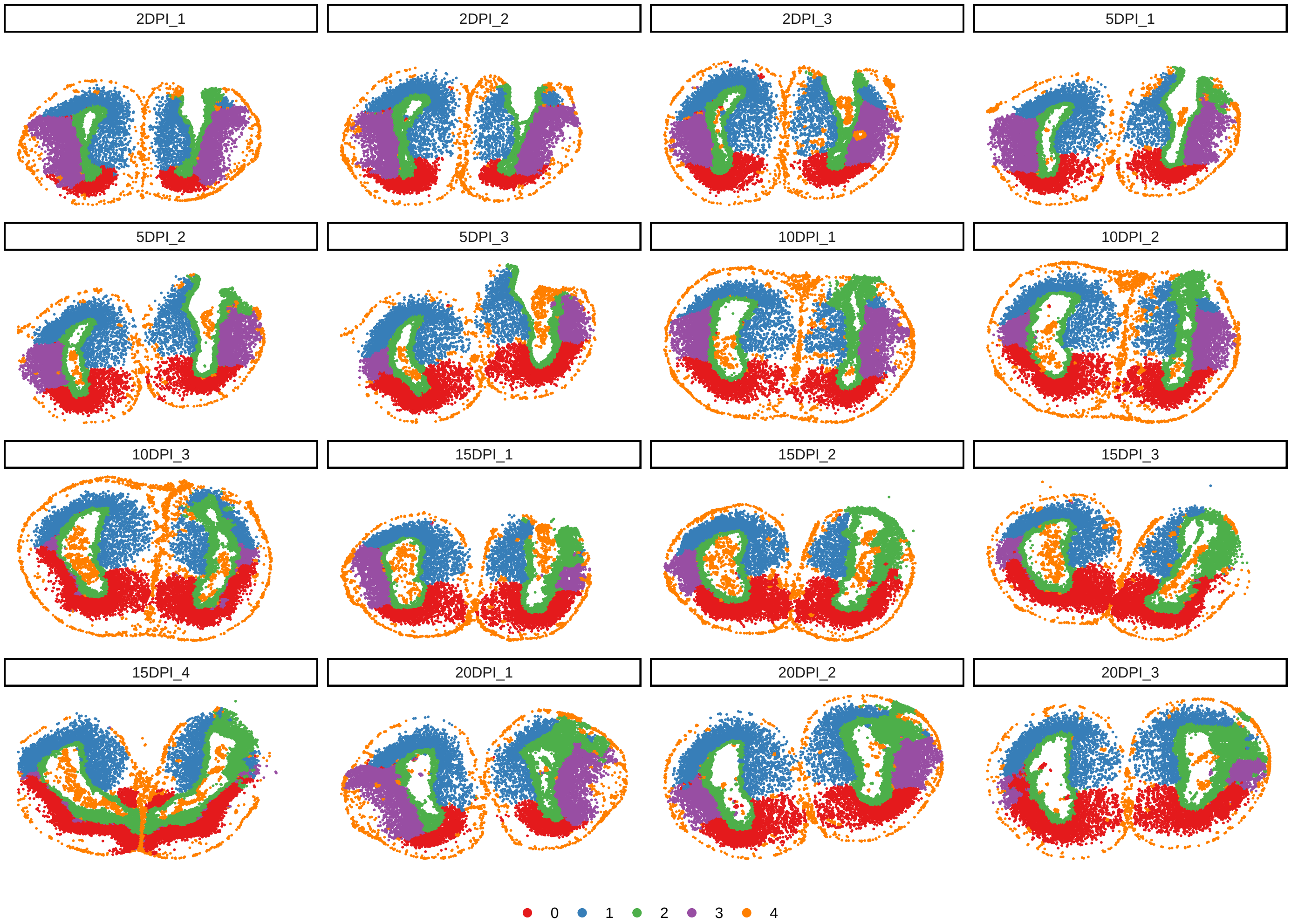


**Supplementary Figure 2:** Spatial domains for the *ARTISTA* dataset, obtained using joint clustering via *Banksy*. These domains were used for both real data analysis and simulation.

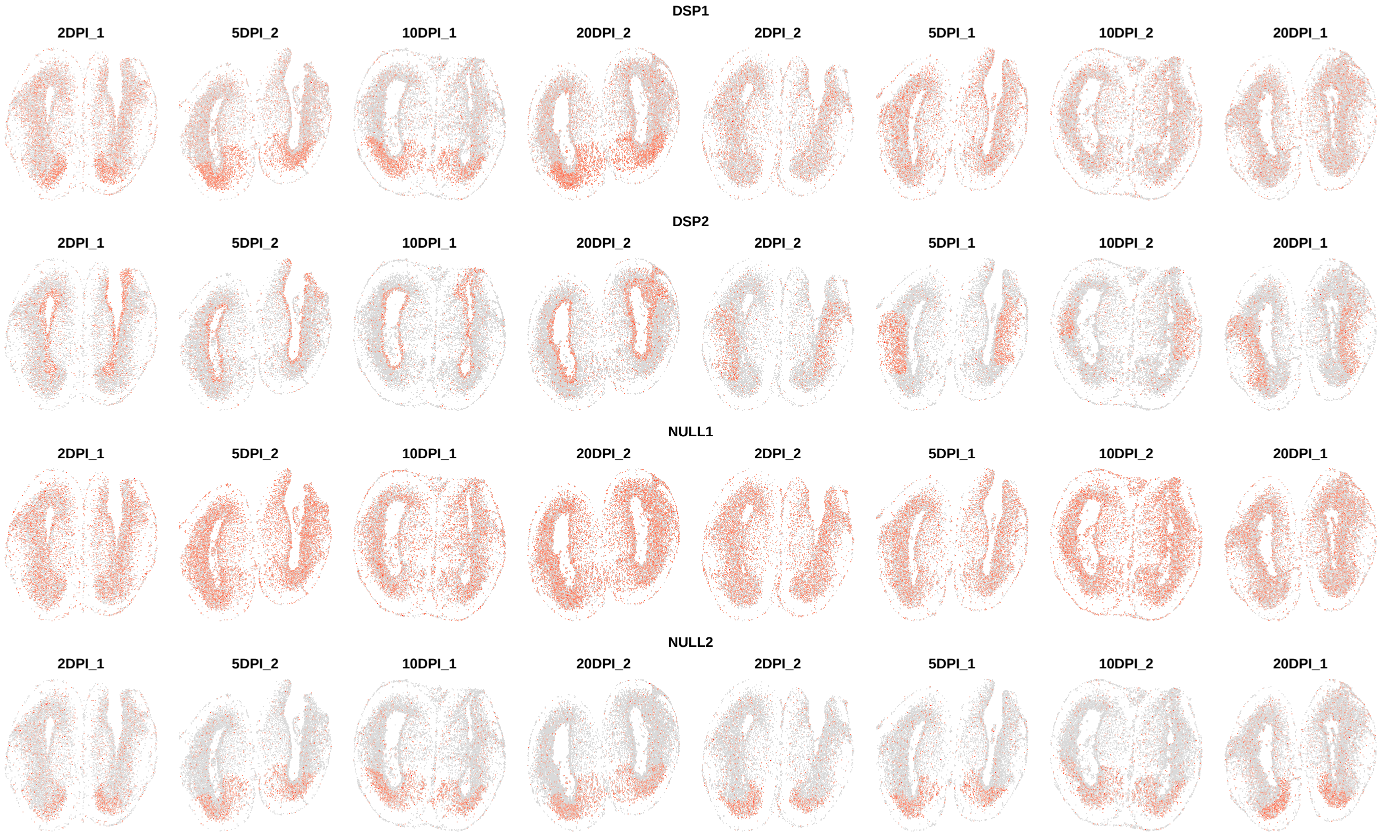


**Supplementary Figure 3:** Examples of simulated DSP genes from the *ARTISTA* dataset, categorized as *DSP1*, *DSP2*, *NULL1*, and *NULL2* (top to bottom), with the selected spatial domain showing higher abundance. The first four columns represent samples from condition 1, and the last four represent condition 2. Cells are colored by expression levels, from light grey (low) to red (high).

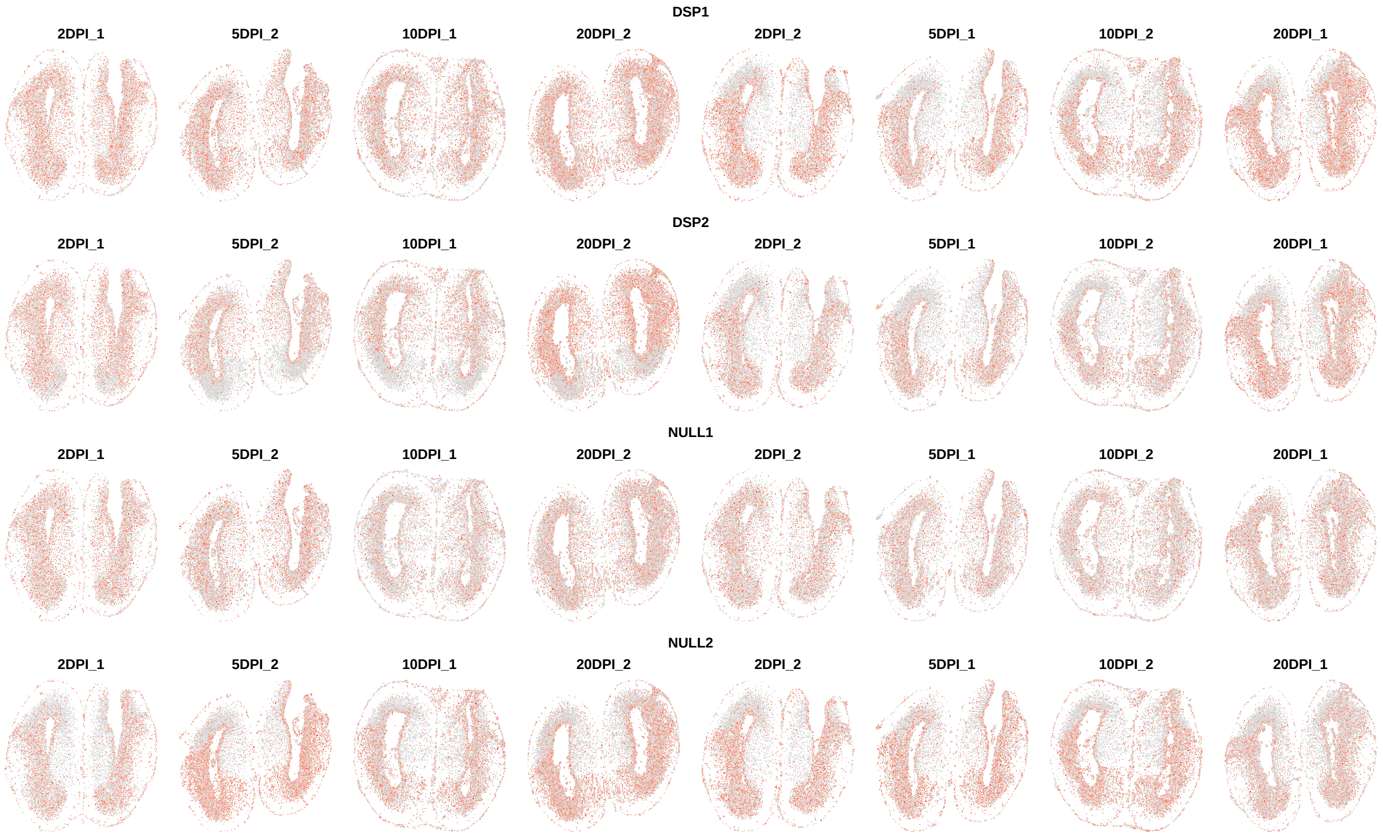


**Supplementary Figure 4:** Examples of simulated DSP genes from the *ARTISTA* dataset, categorized as *DSP1*, *DSP2*, *NULL1*, and *NULL2* (top to bottom), with the selected spatial domain showing lower abundance. The first four columns represent samples from condition 1, and the last four represent condition 2. Cells are colored by expression levels, from light grey (low) to red (high).

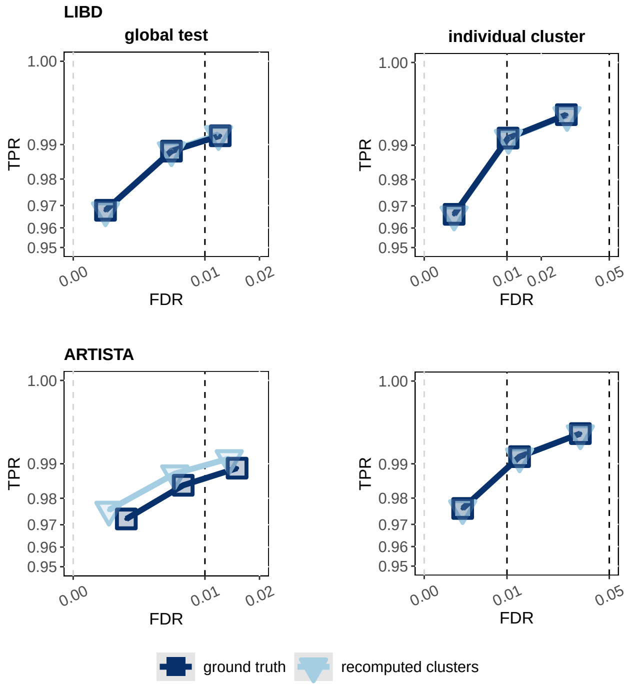


**Supplementary Figure 5:** TPR *vs.* FDR for DSP gene detection in our simulations. Top and bottom rows refer to the *LIBD* and *ARTISTA* data, respectively, while left and right columns indicate the global and individual-domain tests, respectively. Legend indicates the source of spatial clusters used by *DESpace2* with QLF tests: either *ground truth* (annotations used for simulation) or *recomputed clusters* (after simulation via *BayesSpace* for *LIBD* and *Banksy* for *ARTISTA*).

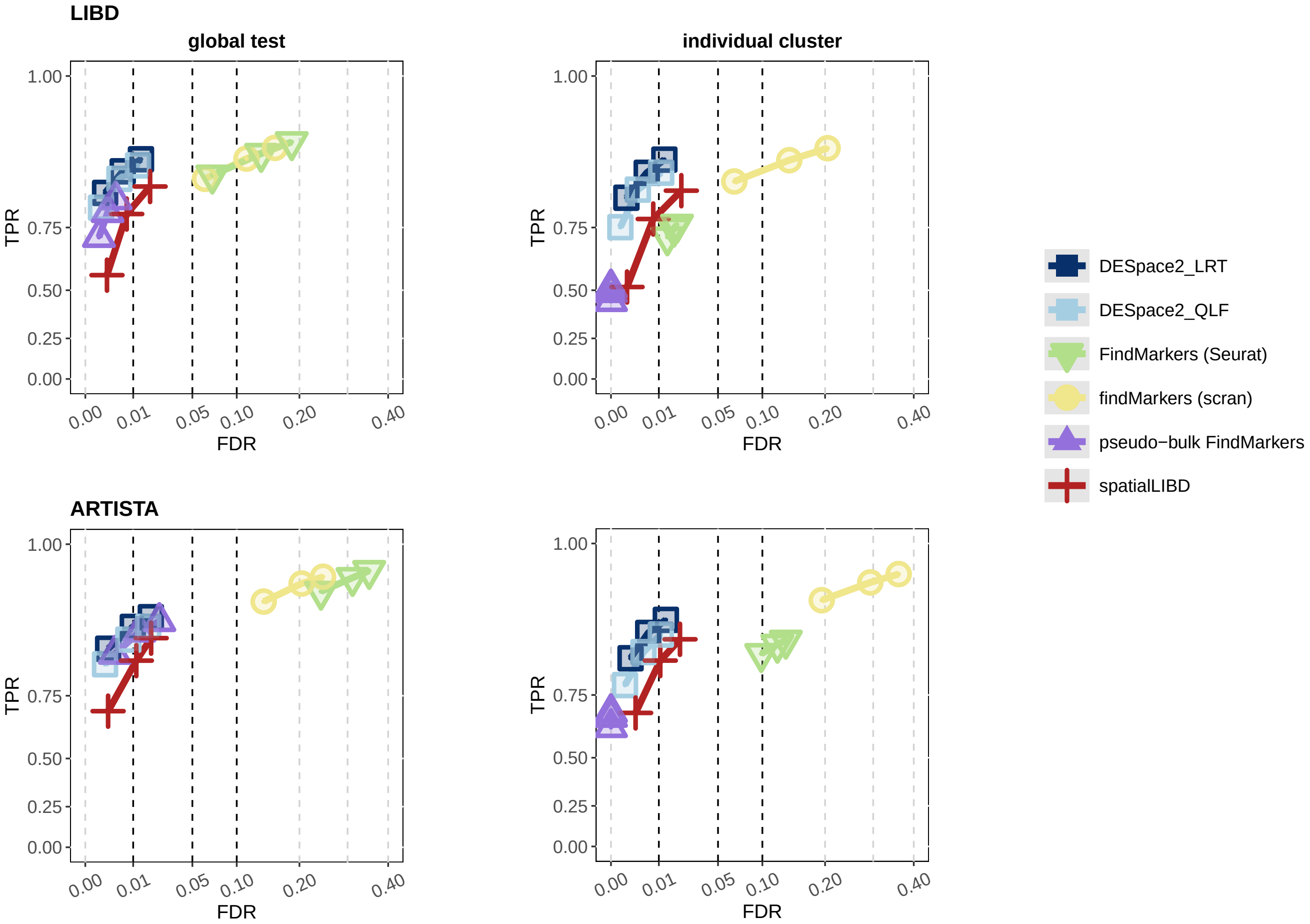


**Supplementary Figure 6:** TPR *vs.* FDR for DSP gene detection in simulations, where the main spatial cluster has lower abundance than the rest of the tissue. Top and bottom rows refer to the *LIBD* and *ARTISTA* data, respectively, while left and right columns indicate the global and individual-domain tests, respectively.

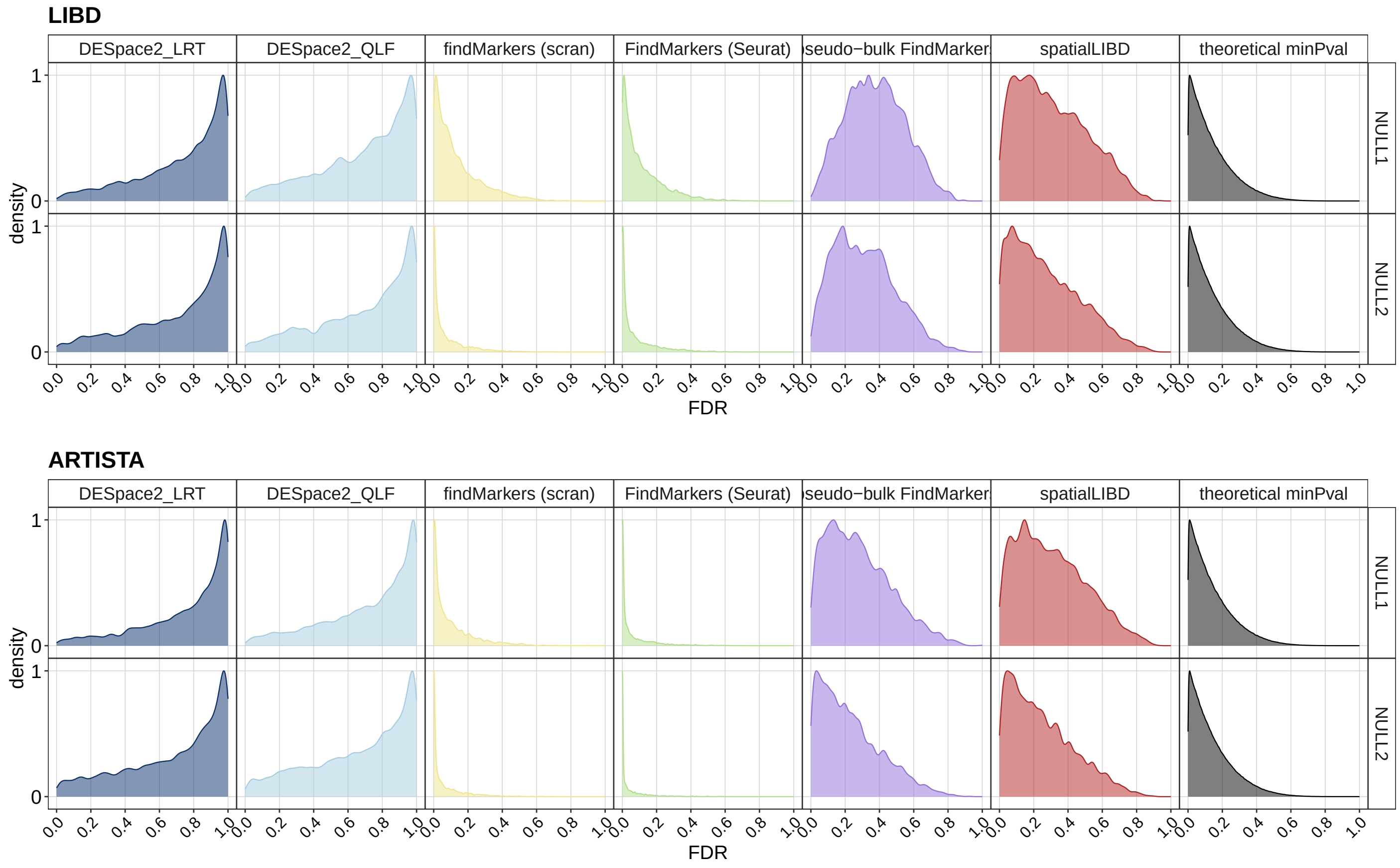


**Supplementary Figure 7:** Density of raw p-values in the null main simulations (i.e., no differences between conditions), based on *LIBD* and *ARTISTA* anchor dataset.

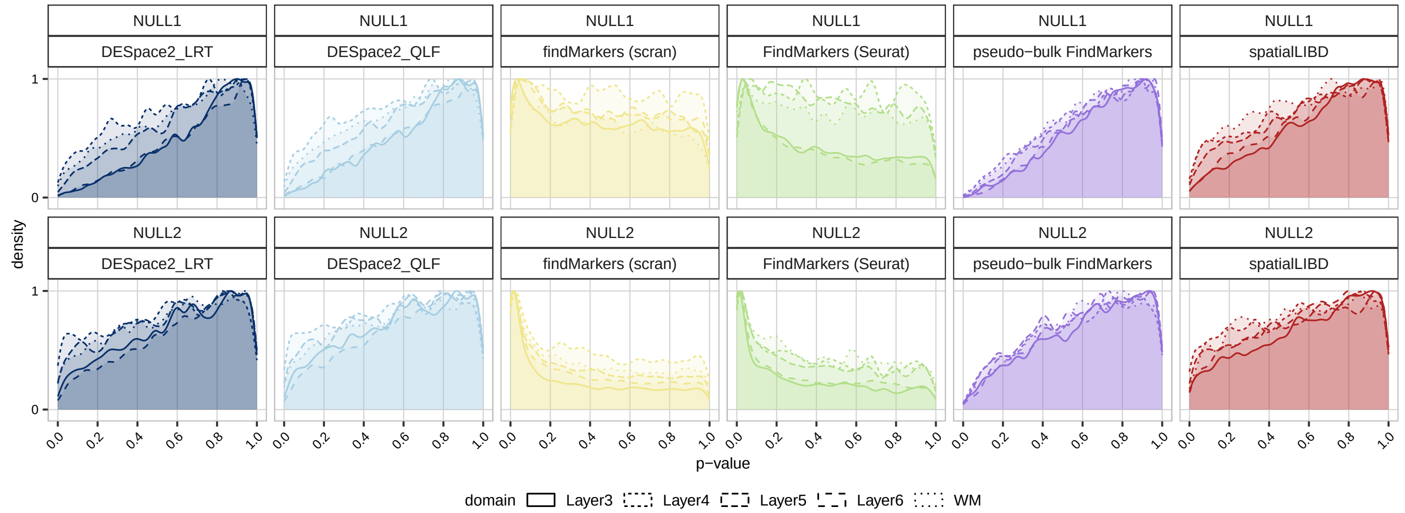


**Supplementary Figure 8:** Density of raw p-values in the null individual cluster simulations (i.e., no differences between groups), based on *LIBD* anchor dataset.

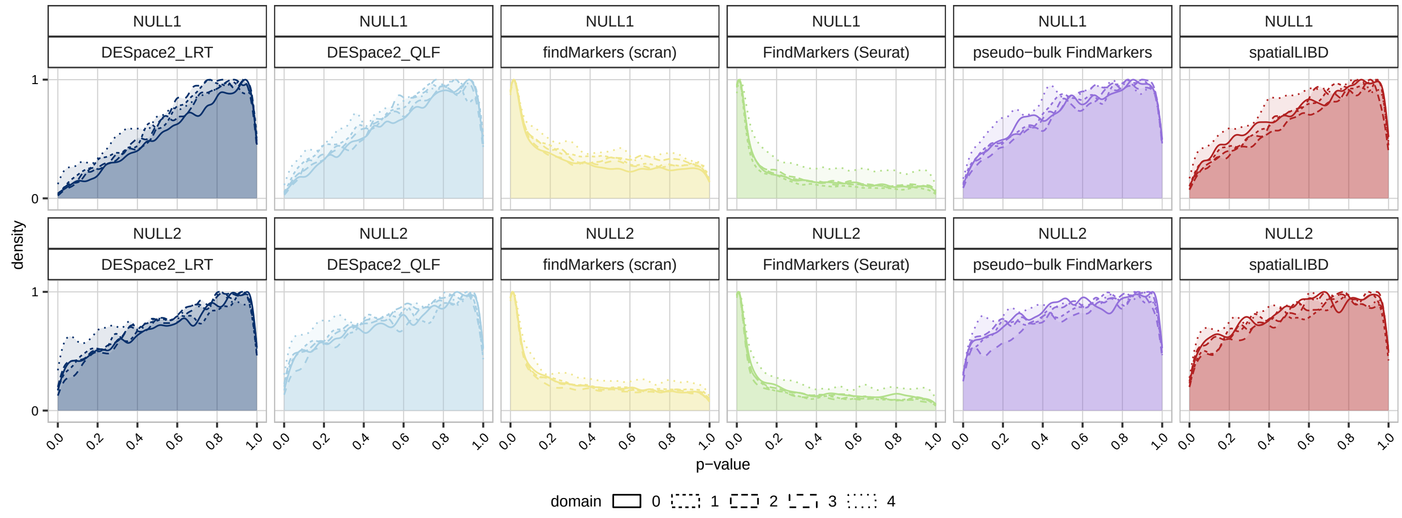


**Supplementary Figure 9:** Density of raw p-values in the null individual cluster simulations (i.e., no differences between groups), based on *ARTISTA* anchor dataset.

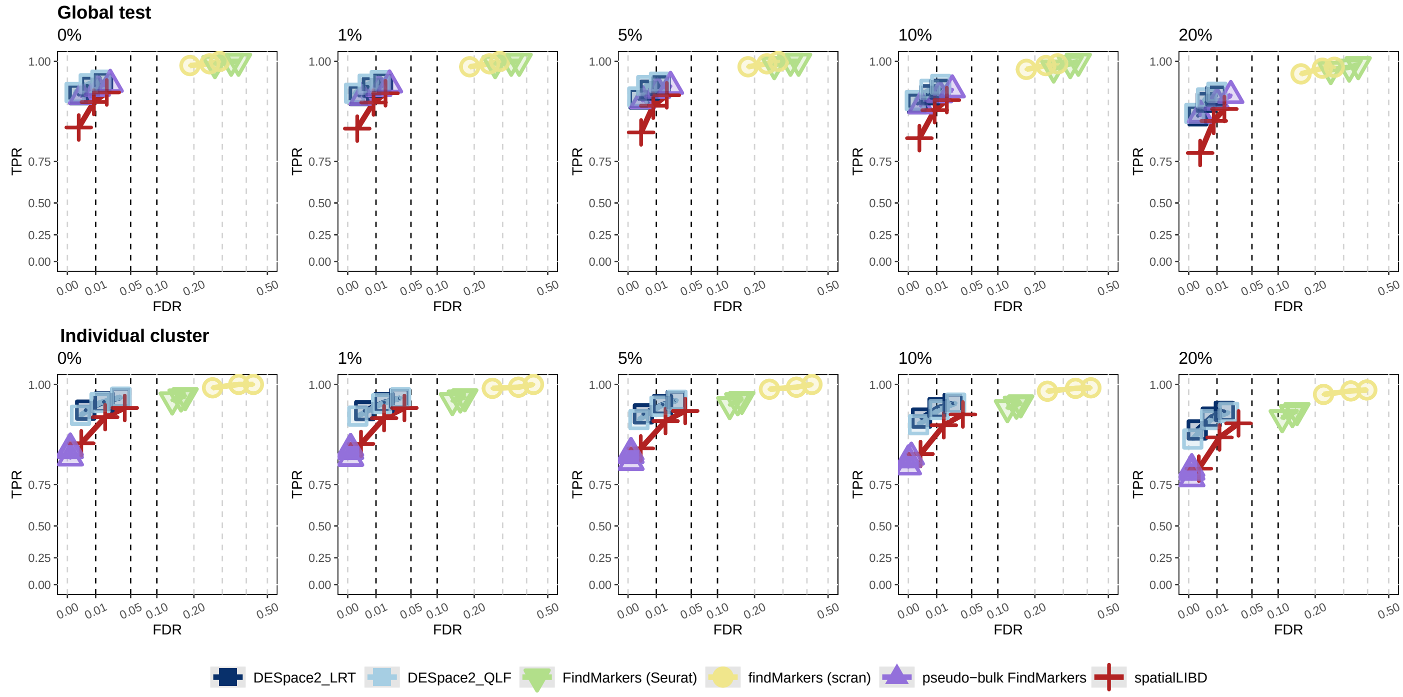


**Supplementary Figure 10:** TPR *vs.* FDR for DSP gene detection in sensitivity analyses using the *ARTISTA* dataset. A subset of cells (0%, 1%, 5%, 10%, and 20%) was randomly reassigned to incorrect clusters to simulate varying levels of mis-clustering. Top and bottom panels refer to the global and individual-domain tests, respectively.

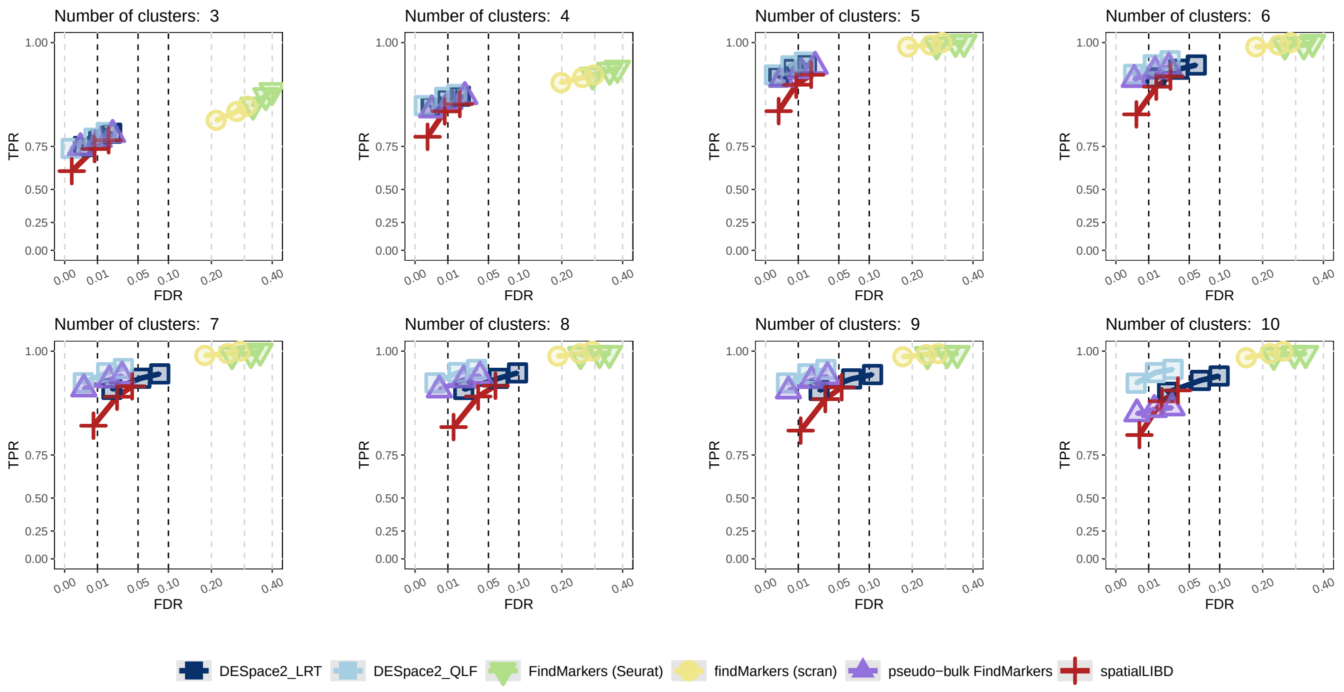


**Supplementary Figure 11:** TPR *vs.* FDR for DSP gene detection in sensitivity analyses using the *ARTISTA* dataset with the global test. Simulated dataset were generated with five clusters, but an incorrect number of clusters (ranging from three to ten) was specified when using *Banksy* clustering.

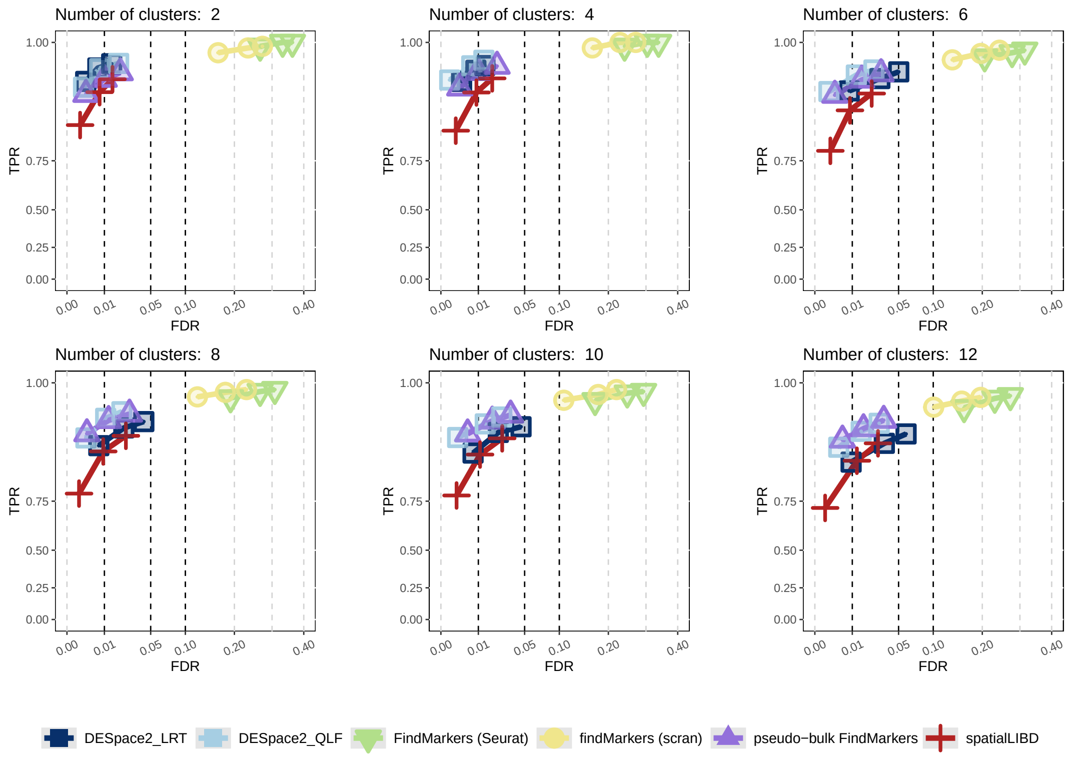


**Supplementary Figure 12:** TPR *vs.* FDR for DSP gene detection in sensitivity analyses using the *ARTISTA* dataset with the global test. Simulated datasets with varying cluster numbers were used to assess robustness to cluster resolution.

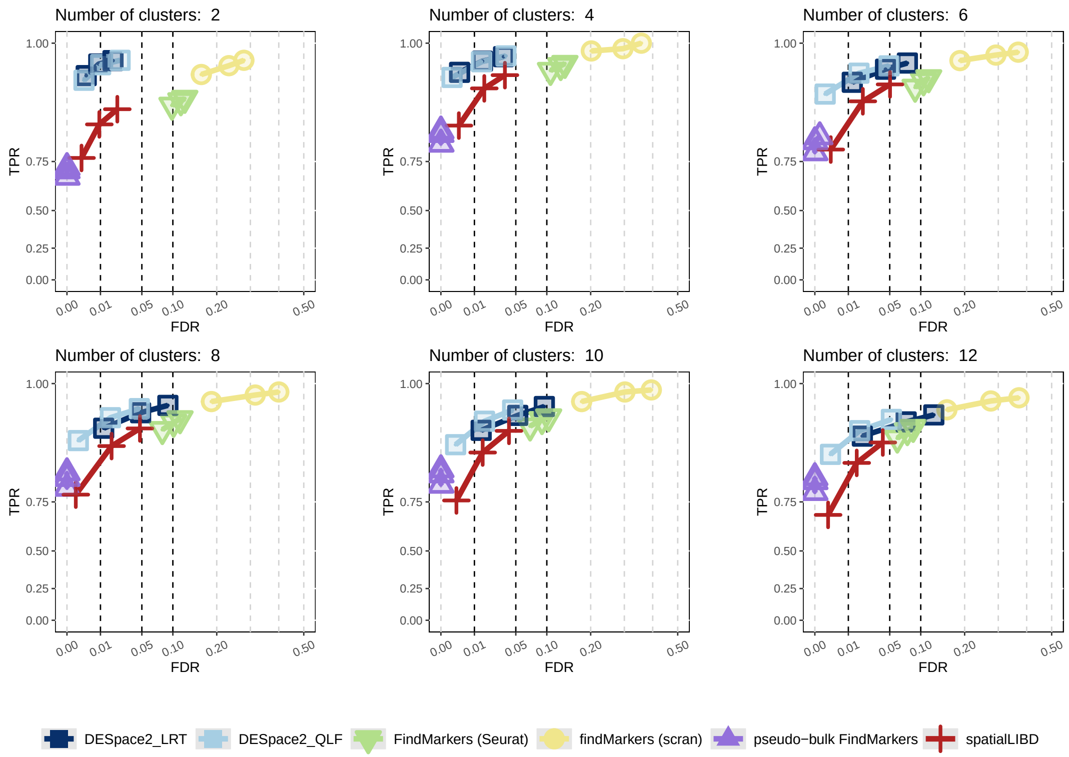


**Supplementary Figure 13:** TPR *vs.* FDR for DSP gene detection in sensitivity analyses using the *ARTISTA* dataset with the individual-domain test. Simulated datasets with varying cluster numbers were used to assess robustness to cluster resolution.

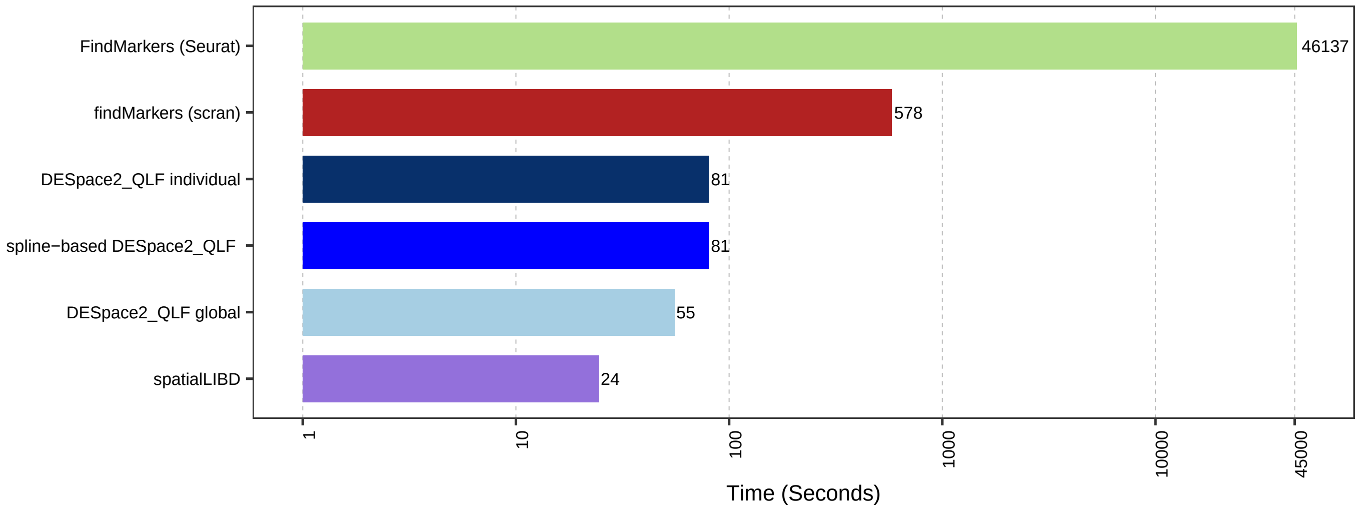


**Supplementary Figure 14:** Runtime (in seconds) for each method on the *ARTISTA* dataset using a 5-condition comparison (2, 5, 10, 15 and 20 days post-injury) across 16 samples.
